# A universal of speech timing: Intonation units form low frequency rhythms and balance cross-linguistic syllable rate variability

**DOI:** 10.1101/2024.12.02.626521

**Authors:** Maya Inbar, Eitan Grossman, Ayelet N. Landau

**Affiliations:** Department of Linguistics, The Hebrew University of Jerusalem, Mount Scopus, 9190501 Jerusalem, Israel; Department of Psychology, The Hebrew University of Jerusalem, Mount Scopus, 9190501 Jerusalem, Israel; Department of Cognitive and Brain Sciences, The Hebrew University of Jerusalem, Mount Scopus, 9190501 Jerusalem, Israel; Department of Experimental Psychology, University College London, London WC1H 0AP, UK

**Author notes:** These authors contributed equally to this work (Eitan Grossman, Ayelet N. Landau).

## Abstract

Intonation units (IUs) are a universal building-block of human speech. They are found cross-linguistically and are tied to important language functions such as the pacing of information in discourse and swift turn-taking. We study the rate of IUs in 48 languages from every continent and from 27 distinct language families. Using a novel analytic method to annotate natural speech recordings, we identify a low-frequency rate of IUs across the sample, with a peak at 0.6 Hz, and little variation between sexes or across the life span. We find that IU rate is only weakly related to speech rate quantified at the syllable level (SR), and crucially, that cross-linguistic variation in IU rate is smaller than cross-linguistic variation in SR. Since SR was shown to be inversely related to information density quantified at the syllable level, we suggest that across languages, IUs are more balanced than syllables in their information content.

## Introduction

What are the foundations of the language system and how are they encoded and realized by our biological machinery? Theorizing about language in a way that is sensitive to the vast diversity of the world’s languages is a challenging endeavor (1), one which raises the question: why have human beings, who share similar brain functions and neurocognitive abilities, developed thousands of distinct languages, each with its unique properties. Research from recent years has introduced exciting ways to tackle this question: first, although the over 7000 distinct languages of the world feature a great degree of diversity, there are robust universal tendencies of linguistic change (2–8). Second, a shifting focus to interactive processes reveals hitherto unknown shared mechanisms across languages (9–11). And finally, considering the unfolding of speech in time has allowed researchers to reconcile apparent diversities across languages. For example, languages differ greatly in their number of distinct syllables, resulting in substantial variation in the amount of information encoded per syllable. Recently, it has been shown that speakers adjust their syllable delivery rate rather consistently within a language, in inverse relation to the amount of information encoded per syllable (12). When syllables are less informative, more are packed into a unit of time and vice versa. In such a way an equilibrium of information transmission per unit time is reached across languages.

The current study is similarly interested in the structure of language in time, and aims to systematically characterize utterance-level prosodic regularities across a genealogically and geographically diverse sample of the world’s languages. Prosody is the domain of language that has to do with how the qualities of our voice, such as pitch and loudness, unfold over time. The qualities of our voice and how they are perceived by our partners in communication have to do with the physical properties of our speech apparatus and of the sound reaching the other’s perceptual apparatus. As such, they are at least partially independent of language-specific linguistic categories. Prosody plays a central role in communication, and – among its many functions (e.g., conveying stance and emotion, initiating action) – is used for segmenting continuous speech into sequences of prosodic phrases. Cross-linguistic research suggests that prosodic phrasing is fundamental to human language and constitutes a universal property of language (13–15). Here we substantiate this claim and characterize the acoustic realization of prosodic phrasing in spontaneous speech in a large and diverse sample of languages. In our work we embrace the close auditory work of generations of speech analysts and phoneticians, and carefully model it acoustically. This body of research has taught us to listen to changes in pitch, delivery rate, and loudness in order to identify units that are termed *intonation units* (henceforth IUs, 13, 16–18), *intonation(al) phrases* (19–22), *tone groups* (23), *elementary discourse units* (24). A characteristic IU is a “stretch of speech uttered under a single coherent pitch contour” (17), with an acceleration-deceleration dynamic of syllable delivery rate and an increase-decrease dynamic of loudness throughout the utterance (14, 25). We touch upon some additional cues that allow us to perceive stretches of speech as separable in later sections.

Our focus on these prosodic phrases stems from their multifaceted importance in communication. For example, at a very basic level, IUs in a given language may become conventionalized and acquire meaning, and become language-specific phonological categories. Yet prosodic phrasing assumes many other roles: it is hypothesized to take on the role of pacing new information in the discourse (26, 27), it allows us to predict in real time when it is our turn to talk (28–31) and it plays crucial roles in language development (32–36). Testing whether IUs assume these roles universally is a fascinating yet complex research program. To carry it out, an important bottleneck needs to be lifted: the reliance on manual annotations of IUs. Therefore, before studying the structure of IUs in time, we begin by devising an automatic annotation option. We then successfully validate its performance compared to traditional manual annotations across languages and use it on a larger corpus of speech recordings.

We study 668 speech recordings of multiple speakers in 48 languages, a sample that represents languages from every continent and from 27 distinct phylogenetic units (i.e., language families). This is one of the largest and most diverse samples to date that tackles questions relating to prosody, and specifically prosodic phrasing. We algorithmically characterize the strength of acoustic boundaries at each and every word in this dataset. This characterization reveals strong correspondences between languages. We show that boundary strength is a dimension clearly dividing the data into two classes, with slight differences between languages. Taking a step further, we show that the resulting classes are directly related to the auditory segmentation of speech into IUs by expert transcribers, for the first time in three languages in addition to English. This validation, we believe, will facilitate and promote studying speech in a manner that is sensitive to its prosodic structure and unfolding over time. And indeed, we rely on our validation work and identify the “acoustic” prosodic phrases across the dataset, which we take as a proxy for IUs.

Finally, recently we and others have shown that IUs succeed each other at a tempo of approximately 1 Hz in several mostly unrelated languages (37, 38). This shared time scale is related to speech production mechanisms, including motor control and biophysical constraints of the speech articulation system (39). In addition, recent neuroscientific findings suggest that low-frequency neural activity contributes to speech production (40) and to the emergence of free thoughts and spontaneous actions (41). In the realm of speech perception and the neural processing involved, low-frequency neural activity is also known to play crucial roles (42–46) and recently we directly related some of this activity to IUs (47). The current study extends the finding that IUs form low-frequency rhythms in spontaneous speech to ever more languages and cultures. Moreover, for the first time, we probe the relationship between syllable rate and IU rate. We show that the two rates are only loosely connected and that languages are more similar to one another in their IU rate compared to syllable rate. This set of results speaks to the question stated at the outset: speakers pace their utterances similarly in time in a wide variety of languages, indeed perhaps universally. The low-frequency temporal structure is a foundation of human language, and it is one that is relatable to the biological mechanisms of our brain and body.

## Results

### Automatically derived IUs show commonalities with manual IU annotations cross-linguistically

We adopt and adapt a method for the hierarchical representation and estimation of prosody, which was previously shown to detect prosodic breaks in English news stories read by 6 speakers (48). In that study, the method reached 85.5% accuracy and 0.73 F1-score relative to expert-made manual annotations of prosodic phrase breaks. Here we applied the method to spontaneous English speech recorded in an informal interview setup (49), and to spontaneous speech in three additional languages: Russian (50), Hebrew (51) and Totoli (52). Table 1 summarizes the method’s performance in prosodic boundary detection in each of these languages. Relative to expert-made manual annotations, the method reached an average of 82.26% accuracy and 0.72 F1-score across the different languages. The average scores indicate moderate classification results in each language to the least, and the average Cohen’s *k* across languages (0.60) is comparable to the agreement between multiple pairs of student annotators in a study surveying the ability to auditorily detect prosodic phrases cross-linguistically (0.58; 14). In addition, in all 34 recordings but one (in English), accuracy was significantly higher than the accuracy expected by simply predicting the most common class in the data. Focusing specifically on English, in which we can compare the method’s performance on spontaneous speech to read speech, the method reached an average of 84.24% accuracy and 0.74 F1-score across the different recordings. Thus, we can conclude that the automatic IU identification pipeline is suitable for cross-linguistic application, and to a spontaneous speaking style.

**Table 1.** Classification results. . Automatic prosodic phrase break detection relative to expert-made manual annotations in four languages.

|  | English | Hebrew | Russian | Totoli |
| --- | --- | --- | --- | --- |
| Precision | 0.76 | 0.90 | 0.71 | 0.90 |
| Recall | 0.73 | 0.47 | 0.59 | 0.89 |
| Specificity | 0.89 | 0.95 | 0.87 | 0.96 |
| **Accuracy** | 84.24% | 73.00% | 77.55% | 94.25% |
| **F1-score** | 0.74 | 0.61 | 0.64 | 0.89 |
| **Cohen's *k*** | 0.62 | 0.44 | 0.48 | 0.85 |

We explore similarities and differences between the manually annotated IUs and the automatically derived IUs (Fig. 1). In Hebrew and in Russian, the automatically derived IUs are longer in the number of words per IU (Fig. 1A). The automatically derived IUs are longer in duration in Hebrew, Russian and Totoli (Fig. 1B). Only in Hebrew were intervening pauses longer between the automatically derived IUs compared to the manually annotated IUs (Fig. 1C). An inspection of several acoustic properties of the automatically derived IUs reveals a large degree of similarity compared to the manually annotated IUs (see *Materials and methods: Acoustic analyses*). The grand average envelope time courses (Fig. 1D), pitch tracks (Fig. 1E) and harmonic ratio time courses (Fig. 1F) do not significantly differ between the methods.

**Fig. 1.**
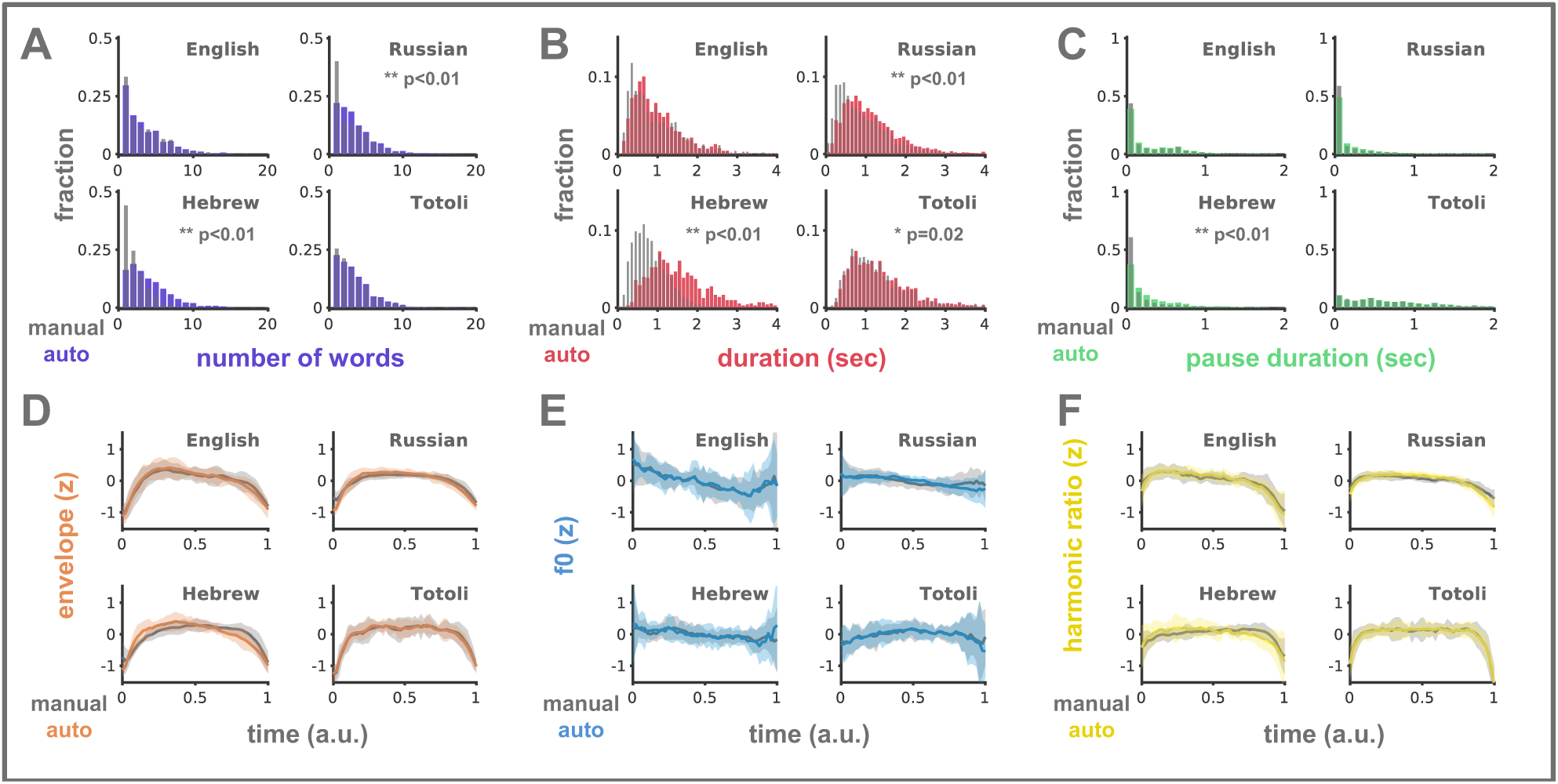
Similarities and differences between the manually annotated IUs and the automatically derived IUs. In each measure, we visualize the data of the automatically derived IUs in color, and in transparent gray the data of the manually annotated IUs. (**A-C**) Probability distributions of the number of words per IU (**A**), of IU duration (**B**), and of pause durations between IUs (**C**). Histogram bins span 0.1 seconds. Wherever a significant difference exists between the two distributions, a p-value is provided in the respective figure. (**D-F**) Time- and scale-normalized time courses of the speech envelope (**D**) f0 (**E**) and harmonic ratio (**F**). Shaded ribbons correspond to the 95% confidence interval corrected for multiple comparisons across timepoints.

With these results in mind, we turn to automatically derive IUs in a large collection of spontaneous speech recordings (53) in 44 additional languages, which did not include IU annotations to date. Inspection of the resulting time- and scale-normalized envelope, f0 and harmonic ratio time courses (Fig. 2A-C) reveals probable correlates of the pattern that lies at the heart of prosodic segmentation, namely, a reset and declination pattern in intensity and pitch throughout the IU, with no significant deviances across languages (see *Materials and methods: Statistical analyses: Deviances across languages in acoustics and IU rate*; see *Supplementary materials: Text S2 and Fig. S1* for analyses at the individual language level). We do not know of a direct demonstration of this pattern in the envelope and harmonic ratio measures, but at least two studies report this pattern in the f0 time course. Specifically, Biron and colleagues (54) show this pitch declination in manually and automatically identified IUs, and directly compare the average f0 values in the time windows 0.15-0.25 and 0.85-0.95. Recently, Ozaki and colleagues (55) similarly demonstrated a notable descending f0 trend in utterances in spoken descriptions across 55 languages. Cohen’s *d* statistics for the difference between the early and late time windows as defined by Biron and colleagues (54) indicate small to medium yet statistically significant average effect sizes across the sample of languages, in each of the measures (envelope: Cohen’s *d*=0.29±0.21 (M±SD), CI [0.23,0.35], p<0.001; pitch: *d*=0.4±0.24 (M±SD), CI [0.33,0.47], p<0.001; harmonic ratio: *d*=0.28±0.16 (M±SD), CI [0.24,0.33], p<0.001). The reset and declination pattern in intensity and pitch is typical of manually annotated IUs (13, 14, 21, 22), and therefore we interpret this result as further validation that the automatically derived IUs can be considered a proxy for IUs, cross-linguistically.

**Fig. 2.**
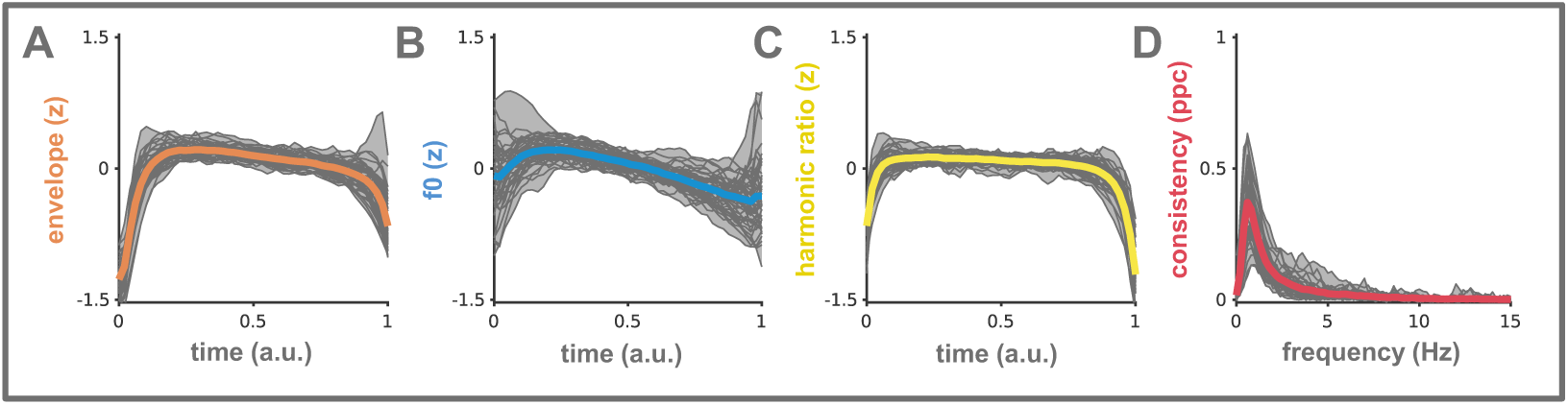
Grand average envelope (A), f0 (B) and harmonic ratio (C) time courses, and phase consistency spectra (D; see next section) of automatically derived IUs. Grand averages across 48 languages appear in color. Grand averages across recordings within a language appear in thin gray lines. Shaded ribbons correspond to 95% bootstrapped confidence intervals that are corrected for multiple comparisons across languages.

### Sequences of IU form low-frequency rhythms cross-linguistically

In a previous study, looking at manually annotated IUs in six languages from around the world, we found that sequences of IUs in spontaneous speech succeed each other at a tempo of approximately 1 unit per second (37). We set out to establish the extent of this effect in spontaneous speech in a broader sample of languages. First, we relied on our original analysis pipeline, which draws on spike-field synchronization studies (56). In our analysis we assume that the speech envelope includes slow periodic modulations. We use these periodic components to estimate the phases of IU onsets, and subsequently examine whether the consistency among phases is higher than expected under the null hypothesis that the slow periodic modulations in the envelope capture word onsets in general, not IU onsets specifically. We find, in all 48 languages, that IU onsets appear at significantly consistent phases of the low-frequency components of the speech envelope, indicating that their rhythms are captured in the speech envelope (Fig. 3). The frequency range of maximal phase consistency in the current sample of languages is 0.6-1 Hz. We next turn to assess whether the spectrum of phase consistency in any language deviates reliably from the grand average across this sample of 48 languages. To this end, we construct a 95% confidence interval around the grand average across languages using a bootstrap procedure, and find that it includes the estimated phase consistency spectra of all languages (Fig. 2D; see *Materials and methods: Statistical analyses: Deviances across languages in acoustics and IU rate*). Therefore, no language can be said to significantly deviate from the others in terms of the alignment of IUs with the periodic components of the speech envelope.

**Fig. 3.**
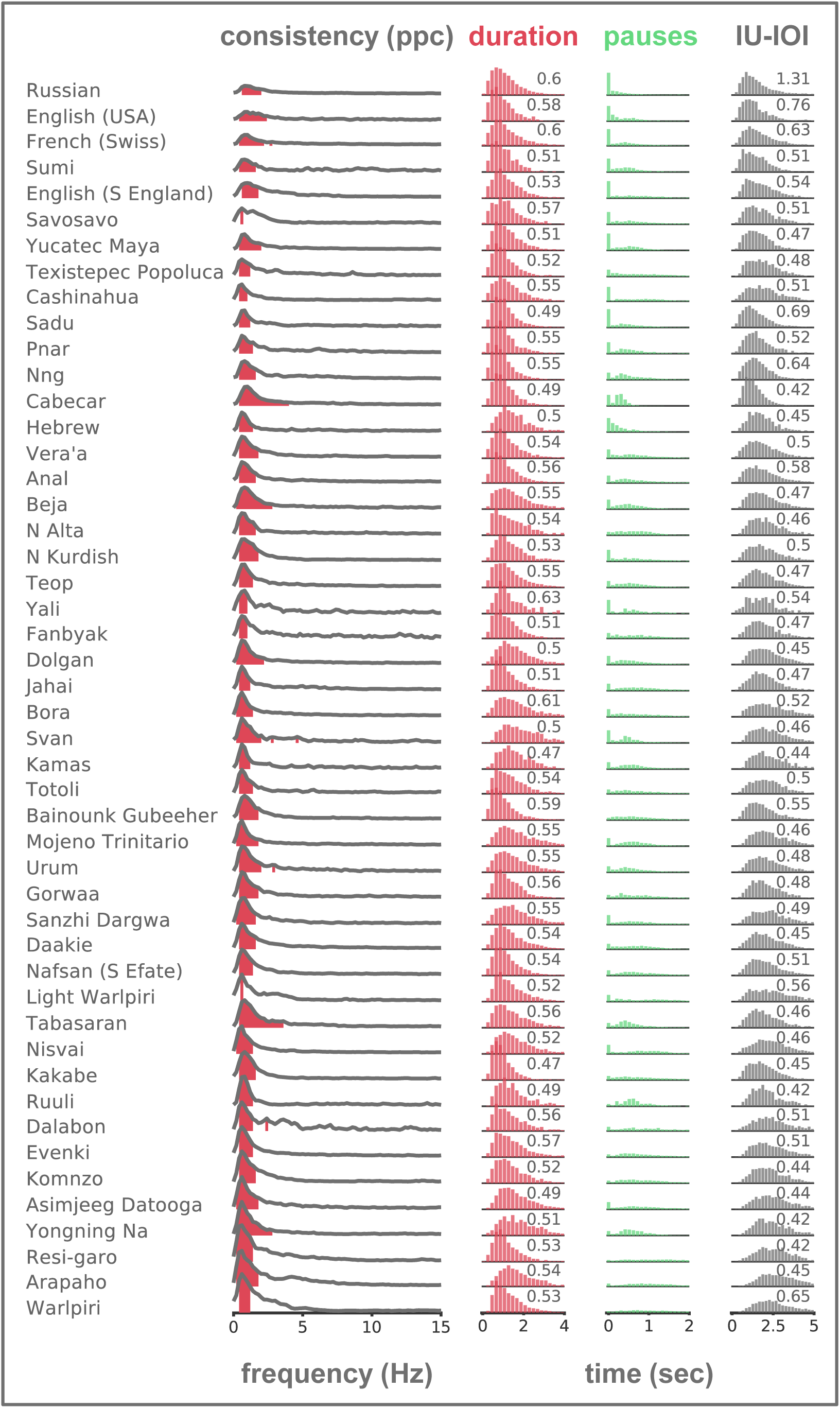
Phase consistency spectra (left) and probability distributions of IU duration (pink), pause duration (green), and IU-IOI values (gray) in 48 languages. Colored regions under the phase consistency spectra correspond to portions of the spectrum that are significantly higher than expected under the null hypothesis, after correction for multiple comparisons across frequencies (see *Materials and methods: Statistical analyses: IU rate*). Languages are sorted in ascending order by their maximal consistency value. The y-axis range of the phase consistency spectra lies between 0-0.63, the maximal consistency value obtained in our sample - in Warlpiri. The y-axis range for the duration and IU-IOI probability distributions lies between 0-0.2 and the histogram bins span 0.2 seconds. The y-axis range for the pause duration probability distribution lies between 0-0.8 and the histogram bins span 0.1 seconds. Coefficient of Variation scores for the duration and IU-IOI measurements appear in the top-right corner of the respective histograms.

We next sought to determine whether the rhythm of IUs characterized above is rooted in the sequence of IUs or rather stems from the durations of IUs themselves. To this end, we calculated per language the Coefficient of Variation (CoV) of the distribution of IU durations (M±SD = 0.53 ±0.03) and the CoV of the distribution of IU inter-onset intervals (IU-IOI values; M±SD = 0.5 ±0.05; see *Materials and methods: Statistical analyses: IU rate*). We tested the paired difference of these measures against zero. On average, IU durations are 4% CI [2%-5%] more dispersed around their mean compared to IU-IOI values. Cohen’s *d* statistic for the difference between two paired samples indicates that this is a medium to large effect size, *d*=0.66. Taken together, these results suggest that rather than producing equally long IUs, speakers tend to balance between IU duration and pauses and in such a way the sequence of IUs gives rise to a rhythm.

### Relation of IU rhythm to syllable-level speech rate

The primary focus of the current paper is the organization of speech into IUs and the temporal structure thereof. However, a broad literature exists that studies speech rate from the perspective of the syllable. As previously mentioned, there is substantial variation between languages in speech rate quantified at the syllable level, and a recent study has found that speakers systematically balance their average syllable delivery rate with language-specific informativeness properties of syllables (12, 57). We sought to connect the two measures of speech rate -- at the IU and syllable levels. First, under the working hypothesis that IUs are related to cognitive mechanisms of attention and memory, we hypothesized that variation across languages in IU rate will be lesser than the systematic variation across languages in syllable delivery rate. In addition, we wished to investigate the possibility that our IU rate results are mediated by effects at the syllable level. To address these points, we explicitly modeled the rate of acoustic landmarks (58, 59), which reflect the onsets of the vocalic nuclei of syllables (henceforth syl-IOI rate; see *Materials and methods: Rate analyses: Syllable rate*), and the rate of IUs (henceforth IU-IOI rate). In both cases, we used a linear mixed-effects approach to account for possible dependencies been measurements, and utilized a model selection procedure to identify the best fitting model.

We found that syl-IOI rate is centered on a mean of 6.77 Hz. This rate is remarkably similar to the mean syllable rate found in previous studies that compared syllable rate in different languages (17 languages in (12), and 55 languages in (55)). Moreover, we replicated a previous finding whereby speaker sex is significantly related to the syl-IOI rate, in the same direction (12). Men have a significantly faster syl-IOI rate of 6.90 Hz. The variation across recordings in mean syl-IOI rate is estimated at 0.35 Hz, and the variation across languages at 0.36 Hz. Nevertheless, the model that includes speaker sex, recording identity, and the language spoken accounts for only a small portion of the overall variance in syl-IOI rate (4%), suggesting that we are missing important information for modeling syl-IOI rate.

Next, we found that IU-IOI rate is centered on a mean of 0.6 Hz, identical to our results based on spectral decomposition (see *Results: Sequences of IUs form low-frequency rhythms cross-linguistically*). The IU-IOI rate is significantly dependent on the local syl-IOI rate, where a faster local syl-IOI rate corresponds to a faster IU-IOI rate. In this model, the total explanatory power is moderate (accounting for 15% of the overall variance in IU-IOI rate), yet the local syl-IOI rate effect accounts for at most 0.8% of the variance in IU-IOI rate. Therefore, it is unlikely that IU rate can be explained away by effects at the syllable level. The variation across recordings in mean IU-IOI rate is estimated at 0.07 Hz, and the variation across languages at 0.12 Hz.

In line with our hypothesis, the models above suggest that variation across languages in mean IU-IOI rate (0.12 Hz) is lower than the variation across languages in mean syl-IOI rate (0.36 Hz). To systematically compare the two, we computed several divergence metrics between IU-IOI estimates of pairs of languages, as well as the same divergence metrics between syl-IOI estimates of pairs of languages (Fig. 4; see *Materials and methods: Statistical analyses: IU rate and syllable rate*). Subsequently, paired permutation t-tests revealed that languages are significantly more like each other in IU-IOI rate in comparison to syl-IOI rate, in all metrics (all p ≤ 0.001).

**Fig. 4.**
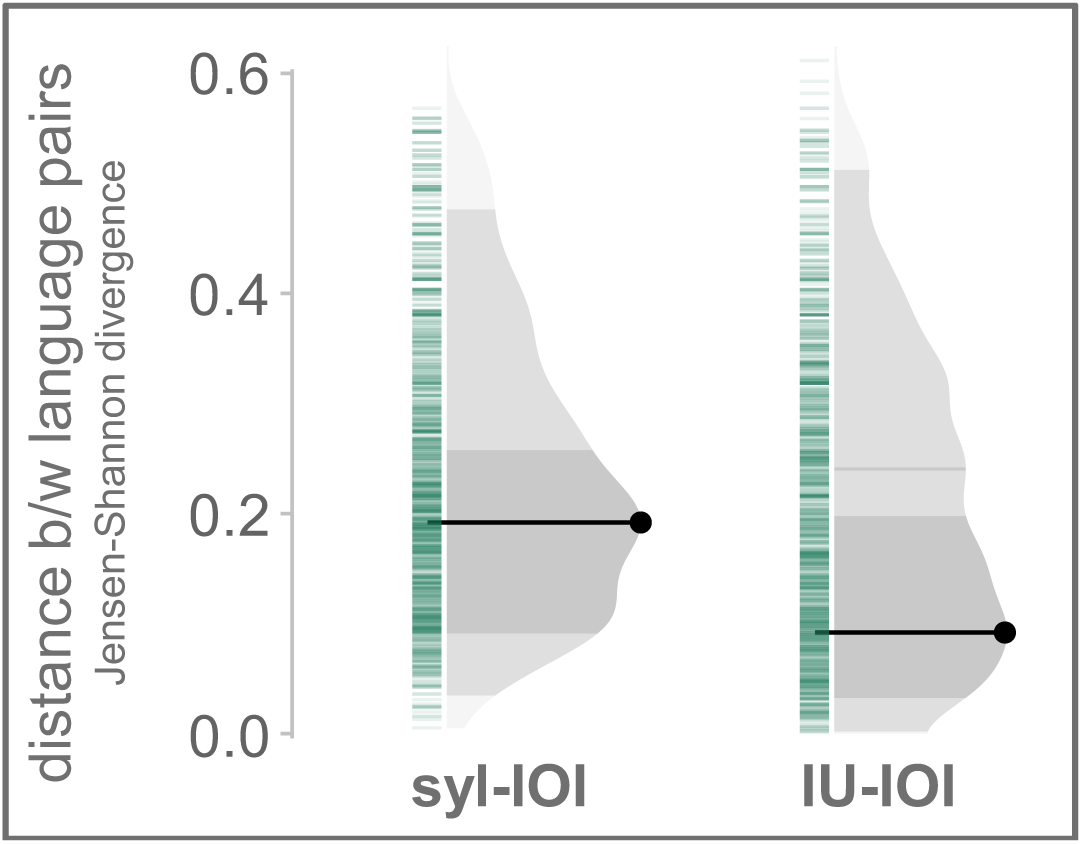
Distances between language pairs in syl-IOI rate and IU-IOI rate, computed using Jensen-Shannon divergence. For each rate measure, green bars indicate individual observations, the black spike indicates the distribution mode and the gray-shaded density plot shows the 50% (dark) and 95% (mid) highest density intervals of the distribution.

## Discussion

In this study, we have investigated prosodic regularities in a diverse sample of 48 languages. We sought to characterize the temporal structure of intonation unit (IU) sequences in spontaneous speech, and to relate our findings to another measure of speech rate. We first validated the performance of an algorithm for automatically identifying IUs cross-linguistically. Then, we revealed a low-frequency rate of IUs across the sample, with a peak at 0.6 Hz, corresponding to an IU beginning every 1.6 seconds. While this rate of IUs varies somewhat between individual recordings and languages, it varies little across speaker sex and age. Finally, we found that syllable-level speech rate (SR) accounts for significant though small variation in IU rate, and more importantly, that the variance across languages in IU rate is significantly smaller than the variance across languages in SR.

Our results support the *universal phonetic-IP hypothesis* (14) by characterizing units with a similar prosodic profile in all the languages of the sample. Language universals are few (1), yet here, through the lens of spontaneous spoken language, we are faced with one. This study therefore demonstrates the importance of considering the temporal unfolding of spontaneous behavior to the study of language and human cognition in general. In future studies, it would be important to investigate the relation of this universal unit to mechanisms of speech production, to bodily rhythms such as breathing, heart rate and eye movements, to mechanisms of attention, temporal prediction, and memory, and to vocalizations in other species.

Recently it has been shown that speakers systematically balance their syllable delivery rate with language-specific informativeness properties of syllables, leading to substantial variation between languages in average SR (12, 57). We show that only a little of this variation affects the delivery rate of a unit hierarchically above the syllable – IUs. This finding suggests that IUs fulfill a role in the pacing of language that is different from the role fulfilled by syllables. Syllables are low-level building blocks that are highly affected by the idiosyncrasies of articulation, while IUs are “planning units”, whose temporal structure relies on mechanisms of attention and memory.

Our findings are consistent with a recent influential neuroscientific model positing that temporally structured brain activity parses and processes incoming speech signals in a hierarchical fashion (60–62). The temporal structure of syllables is relatively well-characterized across the languages of the world, with 4-8 syllables per second (12, 55, 57, 58, 63–65). Here we shed light on a temporal structure at a level higher in the hierarchy, at the level of IUs, which begin every 1.6 seconds. In the brain, activity in this time scale has been especially implicated in studies that are concerned with the question of higher-level speech comprehension, beyond the processing that occurs when we are, for example, listening to backward-played speech, or word-lists (42–44, 66). This suggests that the corresponding linguistic time scale of IUs is especially crucial for speech comprehension and the processing of information. From the perspective of speech production, we know even less about neural mechanisms that might be related to IUs and their temporal structure. A few studies have shown that low-frequency neural activity also contributes to speech production (40) and to the emergence of free thoughts and spontaneous actions (41).

The identification of IUs has been for a long time a laborious manual task, which has limited the breadth of investigations pertaining to IUs, in terms of the languages studied, in the amount of data in each, and across disciplines. We introduce and validate an algorithm to annotate IUs cross-linguistically. We rely on a previous automatic pipeline validated for English (48), and use the same parameters for all studied languages. Here, our sample included only four languages with IU annotations made by expert transcribers, and therefore our ability to tailor parameters to individual languages and investigate the influence of parameter fitting was limited. Future studies might shed light on this matter, and improve the ability to characterize language-specific or even speaker-specific profiles of prosodic phrasing. We see this study as an important step towards studying speech in a manner that is sensitive to its prosodic structure and its unfolding over time. Some of our results are similarly reliant on the automatic identification of syllables cross-linguistically. Future studies should validate whether this method is successful in capturing acoustic landmarks that correspond to the onsets of vowels in languages other than English and Chinese (58), as we assume.

Our study draws a general picture of the temporal structure of IUs in spontaneous production across the world’s languages. The data we analyzed limits our ability to characterize individual variability at the speaker level (the majority of recorded individuals were recorded only once). We are curious as to whether such variability could be traced and explained. Until then, IUs appear to be a universal audio-motor gestalt in human speech, with IUs beginning approximately every 1.6 seconds and with less variance across languages than SR.

## Materials and methods

### Data

#### Unannotated data

We analyzed 634 speech recordings from 44 languages, a sample that represents languages from every continent and from 27 distinct phylogenetic units. This sample was drawn from the Language Documentation Reference Corpus version 1.2 (DoReCo; 53), a database that compiles spoken language corpora originating in documentation efforts of mainly small and endangered languages.

Most of the recordings in the DoReCo database include narratives, either personal or traditional, as well as stimulus retellings, and are recounted by a single speaker at a time. To simplify matters related to acoustic analysis in multi-speaker setting, we limited our sample to the single-speaker recordings (which resulted in the exclusion of data from 3 languages without such recordings). We further excluded data from 4 additional languages whose audio recordings could not be retrieved. While in some cases a single speaker contributed more than one recording, this was not the case for most speakers, and therefore the systematic assessment of individual speaker effects lies beyond the scope of the current study. In addition to audio files, we extracted from the database time-aligned annotations at the word level for each recording. According to the DoReCo annotation conventions, silent pauses were annotated as words, and so we excluded these from our analyses. Table S1 in the Supplementary materials references the specific language corpora used, as well as metadata information about the recordings analyzed in each.

#### Annotated data

For testing and validating the automatic IU identification pipeline, we analyzed 34 speech recordings in 4 languages for which we had expert-made IU transcriptions. Table S2 in the Supplementary materials references the specific language corpora used, as well as metadata information about the recordings analyzed in each. Recordings in three of the languages (Hebrew, Russian, Totoli) had time-aligned annotations at both word- and IU-level, and recordings in one language (English) had no time-aligned annotations whatsoever. We added the levels of time-aligned annotations in English (see *Supplementary materials: Text S1*).

### Automatic prosodic phrase break identification

#### Acoustic boundary strength scores (BS)

Each word in each recording was given a score indicating the strength of acoustic boundary cues at its offset. Scores were calculated using a published algorithm, the Wavelet Prosody Toolkit (48). This algorithm derives fundamental frequency and intensity information from the speech audio signal, and duration information from time-stamped word annotations. As such, the algorithm operates on the signals that correspond to the main cues for an IU boundary. The sum of these signals is decomposed using a continuous wavelet transform in several frequency bands. The algorithm then finds peaks and troughs in the output of the continuous wavelet transform. Based on these peaks and troughs, the algorithm defines for each annotated word a prominence value and a boundary strength value, of which we only used the boundary strength value. In general, the prominence value of a word is defined as the strongest peak within the word. The boundary strength value of a word is defined as the strongest trough between two peaks, namely, between the strongest peak within the word and the strongest peak within the next word. We provided as input to the algorithm the speech audio files and the time-stamped word annotations and applied the algorithm with configurations optimized by Suni and colleagues for detecting prosodic phrase boundaries in English. We set the range of f0 values extracted by the algorithm separately for male (50-350 Hz) and female (100-400 Hz) speakers.

#### Clustering

We utilized the BS scores for the purpose of classifying each word in the data to one of two categories: those that have strong boundary cues and those that have weak boundary cues. The k-means method partitions the BS scores into two groups in a way that minimizes the sum of squares from the datapoints to their assigned cluster centers. We classified words into two groups using the *kmeans* function in the R package *stats* (67). We used the default Hartigan-Wong algorithm with ten starting points.

In compliance with the goal of classifying words into two categories according to their BS scores, we observed that the distribution of BS scores across words tended to be bi-modal (Fig. 5). For the analyses reported below, we classified words into two classes while pooling all words from all speakers across all languages. When considering the entire dataset in this way, the cluster centers were found to be 0.14 and 1.22 (Fig. 5; see black triangles). Based on the identified clusters, we calculated silhouette indices per language to evaluate how similar each point is relative to its own cluster compared to the other cluster (68; Silhouette indices range between ±1, with higher values indicating better separation between clusters and compactness within clusters). The average silhouette width per language ranged between 0.7 to 0.8, which is by convention interpreted as strong clustering. Note that this is the case even though clustering was done while pooling data from all languages in the dataset. An alternative approach would be to classify words into two classes within each language separately (Fig. 5; see pink triangles). The clustering of BS scores into two classes opens several questions for future research regarding cross-linguistic variability. For the current investigation, it lifts the bottleneck of annotating large amounts of data (over 300,000 words). We capitalize on this advance and turn to validate that the pipeline successfully identifies prosodic boundaries cross-linguistically.

**Fig. 5.**
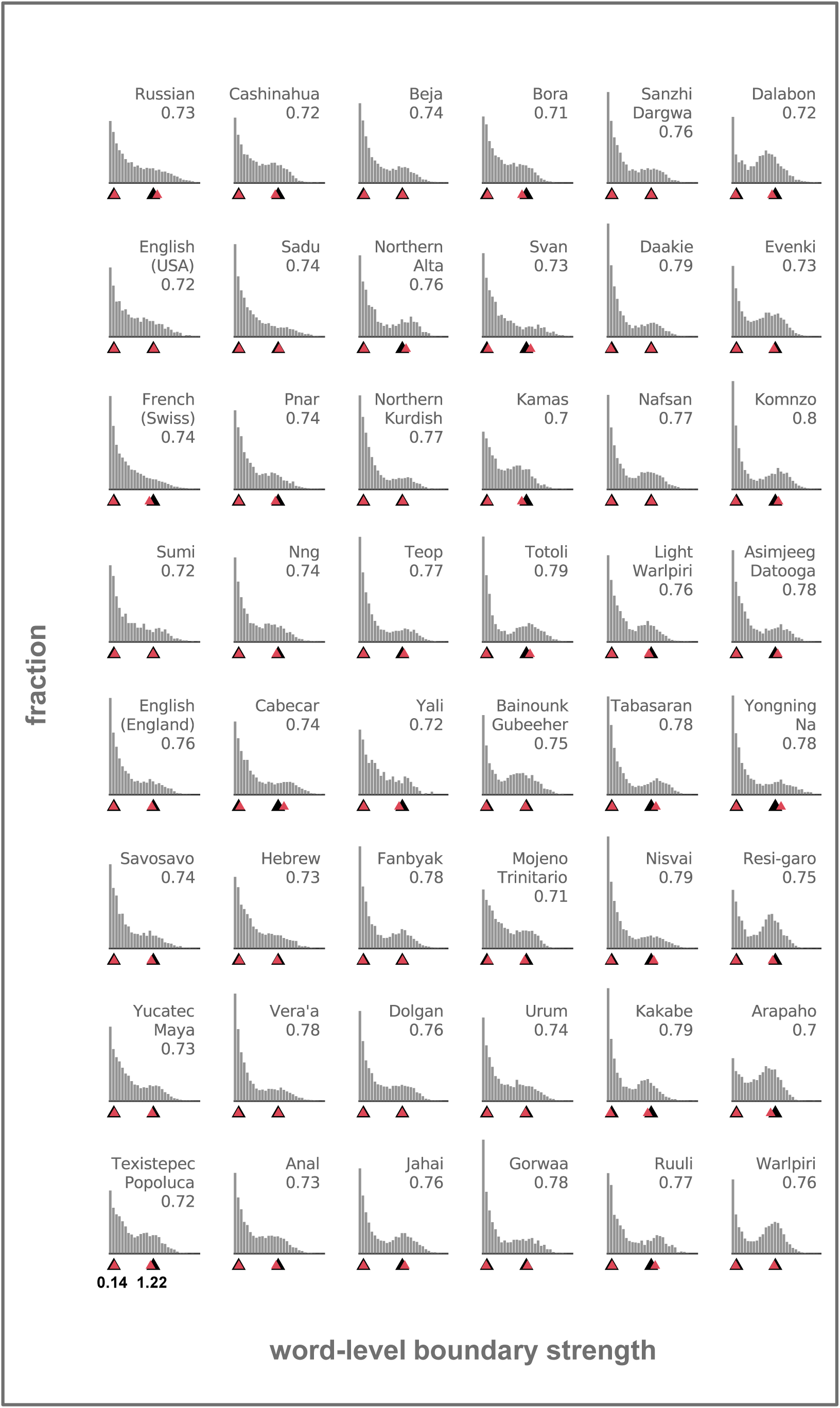
Probability distributions of boundary strength scores. Histogram bins span 0.083 units of boundary strength in the range 0.001 and 2.5, and the y-axis range is identical across languages. The highest histogram bin, in Gorwaa, reaches 0.233. For visualization purposes only, we omitted from the histograms the boundary strength scores that equaled 0, as these were by far the most common (in all languages) and obscured the shape of the histogram. Triangles denote the identified cluster centers, based on the entire pooled data (in black; lower cluster center: 0.14, higher cluster center: 1.22) or at the individual language level (in pink). Average silhouette indices for the clustering based on the pooled data appear below the language label of each histogram.

#### Validation

To assess the validity of the automatic IU identification pipeline, we compared the classification results with expert-made annotations in 4 languages (see *Materials and methods: Data: Annotated data*). To this end, we used the *confusionMatrix* function in the R package caret (69) to compute several metrics: precision (*p*), recall (*r*), specificity, F1-score, and overall accuracy. These measures are derived from true positive (TP), true negative (TN), false positive (FP) and false negative (FN) classification values of IU-final words, and are defined as follows: precision: TP/(TP+FP), recall: TP/(TP+FN), specificity: TN/(FN+TN), F1-score: 2⋅p⋅r/(p+r), and overall accuracy: (TP+TN)/(TP+FP+FN+TN). In addition, we computed Cohen’s *k* (70), a metric for the quantification of agreement over and above chance agreement. To compute these metrics per language we first computed them per recording and then averaged the results across recordings within each language. In addition, per recording, we used a one-sided binomial test with a confidence level of 0.95 to test if the accuracy rate was better than the “no information rate”, the expected accuracy rate when always predicting the most common class in the data.

#### Segmentation

The clustering of BS scores yields a classification of words as either ending an IU or not. From this result we derive a classification of words as either opening an IU or not: we regard words that immediately succeed IU-ending words as beginning a new IU. In this way, and while relying on the annotated onset and offset times of words, we obtain the onset and offset times of IUs.

### Acoustic analyses

We calculated three acoustic signals per recording following the procedures described in the following subsections – the speech envelope, f0 and the harmonic ratio. Subsequently, we epoched each of the three acoustic signals from the onset to the offset of each manually annotated IU in the annotated data (34 recordings in 4 languages), and of each automatically derived IU in the entire sample of recordings (668 recordings in 48 languages). To compare epochs of equal scale and duration we standardized their values relative to the mean and standard deviation of per epoch, and resampled all epochs at 50 equally-spaced time points. Then, per acoustic signal, we computed the average in each recording and the grand average across recordings within a given language.

#### Envelope

We computed the amplitude envelope for each audio file along the following steps: audio files that were recorded in stereo were converted to mono by averaging the channels. Audio files with a sampling rate above 20 kHz were downsampled to 20 kHz. Using the Chimera toolbox, audio files were then band-pass filtered into 9 bands between 100 Hz and half the audio file’s sampling frequency, with cut-off frequencies designed to be equidistant on the human cochlear map (71). Band-pass filtering was performed by using a sixth order Butterworth filter between each two successive cutoff frequencies. Amplitude envelopes for each band (the narrowband envelopes) were computed as absolute values of the Hilbert transform. These narrowband envelopes were downsampled to 1000 Hz and subsequently averaged, yielding the wideband envelope. The wideband envelope was smoothed using a 50 ms sliding Gaussian filter and divided by its maximal value to be on a scale of 0-1.

#### Fundamental frequency (f0)

We computed the pitch track for each audio file using MATLAB’s *HelperPitchTracker* function from the Audio Toolbox, designed to improve pitch estimation of noisy speech signals. Before applying the function, audio files were converted to mono by averaging the channels and downsampled to 8 kHz. The function estimates pitch every 10 ms using partially overlapping windows such that the resulting pitch tracks have a sampling rate of 100 Hz. Multiple pitch candidates are estimated using different pitch detection algorithms, octave-smoothing is applied, and a Hidden Markov Model selects among the candidates based on their confidence and the probability of a certain pitch value moving from one state to another across time in the range 50-400 Hz (we used the pre-trained probability matrix provided by the algorithm). Finally, the function applies a moving median filter with a window length of 30 ms. We set a threshold at 3/4 of the maximal harmonic ratio in the recording (following the conventions in the algorithm) to detect regions that do not include voiced speech and replaced them with NaNs. Finally, we transformed each pitch track to semitones relative to 100 Hz.

#### Harmonic ratio

The harmonic ratio indicates the ratio of energy in the harmonic portion of an audio segment to the total energy in that segment. The ability to extract the fundamental frequency depends on whether periodicities can be reliably extracted from the signal, and this is why the MATLAB function *HelperPitchTracker* relies on the harmonic ratio. However, the dynamics of voicing during speech are interesting in their own right. The function *harmonicRatio* from MATLAB’s Audio Toolbox is called from within *HelperPitchTracker*, and we record its output to inspect its dynamics relative to IU boundaries in addition to the dynamics of pitch.

### Rate analyses

#### IU rate

We analyzed the relation between IU onsets and the speech envelope using a point-field synchronization measure. In this analysis, the rhythmicity of IU sequences is measured through the phase consistency of their onsets with respect to the periodic components of the speech envelope. Using this method, we have previously shown that IU onsets appear at significantly consistent phases of the low-frequency components of the speech envelope in six languages (37). We extracted 5 second windows of the speech envelope centered on each IU onset, and decomposed them between 0-15 Hz using Fast Fourier Transform (FFT) with a single Hann window, no padding or smoothing across frequencies, and following demeaning. This yielded phase estimations for frequency components at a resolution of 0.2 Hz. We then measured the consistency in phase of each FFT frequency component across speech segments using the pairwise-phase consistency metric (PPC; 56), yielding a consistency spectrum. We computed a PPC spectrum per recording and language, and averaged the spectra across recordings within a language. This method is adopted from the study of rhythmic synchronization of neural spiking activity and Local Field Potentials.

As an additional measure of IU rate, we calculated the intervals between IU onsets (inter onset interval, and henceforth, IU-IOI) in each recording and language, from which we calculate the IU-IOI rate. The unit of IOI is seconds, and IOI rate is the reciprocal of IOI and is measured in Hertz. To aid the convergence of models in subsequent statistical analyses we excluded IU-IOI rates that were above or below thresholds calculated per individual recording. Given IQR, the range between the first (Q1) and the third (Q3) quartiles of IU-IOI measurements in a recording, the lower threshold was defined as Q1-3·IQR and the upper threshold was defined as Q3+3·IQR. To this end, we used the *identify_outliers* function in the R package *rstatix* (72) and the *anti_join* function in the package *dplyr* (73). This resulted in excluding 1.04% of the 91540 IU-IOI rates.

#### Syllable rate

While the preceding section dealt with the rate of IUs, in the following we turn to discuss our methods for estimating speech rate from the perspective of the syllable, with the aim of studying its relation to our main findings on the rate of IUs.

A precise estimation of speech rate at the syllable level requires fine-grained transcription and detailed knowledge of each language’s phonological system (see Text S2 in 12, 55). An alternative is to estimate speech rate through automatically identifying syllable nuclei (usually vowels) via acoustic analysis. However, we are somewhat lacking in evidence on the robustness of automatic methods in estimating speech rate, if not the contrary – that they might systematically err in performance in different languages. For example, Coupé and colleagues (12) considered such an algorithm yet deemed it not sufficiently reliable, as did additional studies that considered large samples of languages (55, 74). Since then, however, another approach for detecting vowel onsets automatically was reintroduced in the literature (59), and in a recent comprehensive assessment it was found to outperform other detection methods in two unrelated languages and diverse speaking contexts (58), even if not perfectly replicating gold-standard inter-vowel time series. Of course, much work is still required to assess the robustness of this method across languages, yet we chose nonetheless to rely on it here for considerations of time and the scope of this study.

We utilized the scripts published by MacIntyre and colleagues (58) to estimate syllable-level speech rate in our dataset (henceforth, SR). These scripts consider 5 different procedures to extract envelopes along with 5 event detection methods (such as peaks in the envelope or peaks in the envelope’s first derivative). Of the 25 possible acoustic landmarks, we a-priori chose the one method that was found by MacIntyre and colleagues (58) to be the most robust, termed “envelope3/peaks in the first derivative”. This is the precise acoustic landmark used by Oganian & Chang (59), and which they found to be specifically encoded by a defined region in the Superior Temporal Gyrus during speech perception. Unlike MacIntyre and colleagues (58), we applied the algorithm to the continuous speech recordings (unsegmented in any way) rather than to windowed data, to avoid conflating in unknown ways the acoustic landmark detection with the segmentation of spontaneous speech currently under discussion. However, to ensure that the resulting SR will not be calculated over periods of silence, we subsequently used the onset and offset times of IUs (see *Materials and methods: Automatic prosodic phrase break identification: Segmentation*) to epoch the continuous time course of acoustic landmarks, thus excluding any non-speech acoustic landmarks that might have been calculated by mistake. We then calculated the intervals between acoustic landmarks within each epoch (an epoch with *n* detected acoustic landmarks contributed to the calculation of *n-1* intervals, meaning that an epoch had to contain at least two acoustic landmarks in order for an interval to be calculated). For simplicity, we refer to these measurements as syl-IOI, yet we remind that acoustic landmarks more precisely reflect the onsets of the vocalic nuclei of syllables. From the syl-IOI we calculate the syl-IOI rate. The unit of IOI is seconds, and IOI rate is the reciprocal of IOI and is measured in Hertz. To aid the convergence of models in subsequent statistical analyses we checked if any syl-IOI rates were above Q3+3·IQR and below Q1-3·IQR (calculating quartiles per individual recording), with the aid of the *identify_outliers* function in the R package *rstatix* (72) and the *anti_join* function in the package *dplyr* (73). None of the syl-IOI rates exceeded these criteria. Finally, to reach an IU-by-IU measure of SR, we either averaged the syl-IOI rates within each IU epoch, simply recorded the syl-IOI rate if there was only one, or recorded NA if no syl-IOI was detected in it.

### Statistical analyses

#### Validation

##### Number of words, duration and intervening pauses

To compare these measures in the automatically derived IUs and in the manually annotated IUs we conducted a series of nonparametric statistical tests per measure and language. The test statistic was the estimate for the difference between the two methods in the measured variable. The estimate was obtained in a mixed-effect general linear model of the measure against the segmentation method, with a by-file random intercept (75). We created a permutation distribution for each statistic by shuffling segmentation method labels 100 times. This permutation distribution represents the expected difference between the segmentation methods under the null hypothesis, as it breaks the association between measure and method. This procedure created 3(measures) x 4(languages) = 12 permutation distributions which we compared to 12 observed test statistics, namely, the estimated difference in each of the measures and in each of the languages between the manual and automatic IUs. As a p-value for each test, we calculated the proportion of times that the permutation procedure resulted in an estimate larger than the observed estimate for the difference between the manually and automatically derived IUs.

##### Envelope, f0 and harmonic ratio

To compare these measures in the automatically derived IUs and in the manually annotated IUs we computed confidence intervals around the grand average time course in each measure and language. We first computed the mean and standard error, and then multiplied the standard error by the critical t value. The critical t value was calculated with degrees of freedom according to the number of non-NaN observations in each timepoint, and for an alpha level of 0.025/50=0.0005, to correct for multiple comparisons across the different timepoints via Bonferroni correction. The intervals overlapped in all measures and languages.

#### IU rate

We assessed the statistical significance of peaks in the average PPC spectra using a randomization procedure. Per language, we created 100 sets of surrogate average spectra. These surrogate spectra were calculated using the speech envelope as before, but with temporally permuted onsets that maintained the association with word onsets (i.e., we sampled onsets from among all words in each recording without replacement). Each iteration resulted in an average phase consistency value at each frequency. However, we generated a single randomization distribution from this procedure by selecting, per iteration, the maximal consistency value across frequencies. We then calculated, for each frequency, whether the observed consistency value exceeds the 95% percentile of the randomization distribution of maximal statistics. This approach corrects for multiple comparisons over the different frequencies (76).

In an additional analysis, we tested whether the Coefficient of Variation (CoV) of the distribution of inter IU onset intervals (IU-IOI) was different from the CoV of the distribution of IU durations. To compute the CoV of each distribution per language we first excluded observations (IU-IOI and IU duration) exceeding ±2.5 standard deviations from the mean within each language. We additionally excluded the entire data from two languages, Russian and American English (from the validation set), as this data comes from conversational corpora and therefore a substantial number of IU-IOIs are conflated by the occurrence of speaker change. We then subtracted per language the CoV of the IU-IOI distribution from the CoV of the duration distribution, and averaged across the 46 remaining languages to obtain the observed test statistic. We used a nonparametric bootstrap procedure with 100,000 iterations to construct a confidence interval around this statistic. On each iteration, we sampled with replacement 46 CoV differences and averaged the sample. This creates a distribution of average CoV differences that could have been obtained had the sample of languages been composed slightly differently. We computed the limits of the confidence interval as the 2.5% and 97.5% percentiles of this distribution. If this interval does not include 0, we can conclude that 0 is an unlikely value for the difference between the CoV of the IU-IOI distribution and the CoV of the duration distribution.

#### Deviances across languages in acoustics and IU rate

We assessed whether individual languages differed in their envelope, f0 and harmonic ratio time courses and PPC spectra from the grand average time course (or spectrum, in the case of the PPC measure) across languages. To this end, we constructed a confidence interval using a nonparametric bootstrap procedure. On each iteration, we sampled with replacement time courses (or spectra) per language, according to the number of languages in our sample. Each iteration resulted in an average value per timepoint (or frequency) and language. However, we generated a single distribution per timepoint (or frequency) by selecting the maximal and minimal values across individual languages. We computed the limits of the confidence interval per timepoint (or frequency) as the 2.5% percentile of the distribution of minimal values in that timepoint and the 97.5% percentile of the distribution of maximal values. We then calculated, for each language, whether the observed value in a given timepoint (or frequency) exceeds this interval. This approach corrects for multiple comparisons across the different languages (76). We do not additionally correct for multiple comparisons across time bins as there were no significant effects following the comparison across languages.

#### IU rate and syllable rate

We followed the approach in Coupé and colleagues (12) and conducted several statistical analyses:

(1) We modeled SR using the automatically derived syl-IOI rate.
(2) We modeled the IU rate using the IU-IOI rate; in both these cases, we used a linear mixed-effects approach, to account for possible dependencies been intervals originating from the same recording, language, language family, area, age and sex. In the modeling of IU-IOI rate we additionally considered the local, IU-level SR as a potential source of variance in the distribution of IU-IOI rates.
(3) We compare the divergence between languages in IU rate and in SR, and show that it is significantly lower in the case of the IU rate.

As a preliminary analysis, we fitted all unique combinations of the predictor variables mentioned above (127 combinations), separately for the dependent variables syl-IOI rate and IU-IOI rate. We used the BIC to select the best fitting model, thus balancing the likelihood with the number of model parameters to avoid overfitting.

In both cases, the selected models included the same 2/4 random intercepts we considered (recording and language, but not family and area). Unlike the case in Coupé and colleagues (12), our speech material does not include multiple texts of the same kind (each recording featured unique material) and does not include multiple recordings for most speakers. Therefore, we could not test these effects, but we used random intercepts per recording instead. In addition, our sample usually does not include more than one language per family, and this is probably a source of divergence in our selected syl-IOI rate model compared to Coupé and colleagues (12; they too note that family had a very small effect beyond language). Finally, while the diverse data at hand allowed us to estimate random intercepts per linguistic area, the models including them had a lower BIC compared to the selected model. The selected syl-IOI rate model included only a fixed effect for speaker sex, identical to Coupé and colleagues (12). The selected IU-IOI rate model included only a fixed effect for IU-level SR (see *Materials and methods: Main analyses: Speech rate*).

We fitted the models using the R package *lme4* (75), and tested the significance of the fixed effects using the package *lmerTest* (77). Speaker age and IU-level SR variables were standardized, such that their estimated coefficients could be interpreted as the expected change in IOI for a standard deviation change in age or IU-level SR, and the intercept as the expected IOI for an average age and IU-level SR. The speaker sex variable was coded as a sum contrast so that the estimated intercept could be interpreted as the average across sexes.

Inspecting the results of the selected models, the total explanatory power of the syl-IOI rate model is weak (conditional *R^2^* = 0.04) and the part related to the fixed effects alone (marginal *R^2^*) is 0.0026. The intercept of the model, corresponding to the average syl-IOI is at 6.77 (95% CI [6.66, 6.88]). Speaker sex is significantly related to the syl-IOI rate, with men having significantly higher syl-IOI rates (*β* = 0.13, 95% CI [0.10, 0.17], t(572937) = 7.80, p < .001).

The total explanatory power of the IU-IOI rate model is moderate (conditional *R^2^* = 0.15) and the part related to the fixed effects alone (marginal *R^2^*) is 0.0085. The intercept of the model, corresponding to the average IU-IOI rate with average syl-IOI rate, is at 0.60 (95% CI [0.57, 0.64]). IU-level syl-IOI rate is significantly related to the IU-IOI rate, with faster rates of syl-IOI associated with faster rates of IU-IOI (*β* = 0.03, 95% CI [0.03, 0.04], t(86299) = 27.43, p < .001).

Finally, we extracted from the selected models the estimated syl-IOI and IU-IOI rates per recording for the following final analysis. In this analysis, we followed precisely the steps and code in Coupé and colleagues (12) for inspecting pairwise distances between languages, except we consider the divergence between languages in syl-IOI and IU-IOI rates. For each of the two variables, we computed all the possible 48·(48–1)/2 = 1128 unique pairs of languages, and compared the two languages’ distributions of a given IOI rate variable. As in (12), we used five methods for a robust comparison (Hellinger distance, Jensen-Shannon divergence, Kolmogorov-Smirnov distance, Kullback-Leibler divergence, and chi-square divergence) with the aid of the *distance* function from the R package *philentropy* (78; all divergence methods provided equivalent conclusions). We subsequently tested paired differences between corresponding distance metrics for syl-IOI rate and IU-IOI rate across the sample of languages using permutation t tests with 1000 permutations.

## Acknowledgments

We thank Anat Perry for access to the Hebrew data we report on, Shlomi Frige for help with its annotation, and Nadav Matalon, Amit Avigdor, Hagai Berenson, and members of the Brain, Attention and Time lab for discussion. We especially thank Frank Seifart and Nikolaus Himmelmann for the resources and thoughts they shared with us. Our work would not have been possible without them and without the substantial efforts of language documenters and the people they record all over the world.

## Author contributions

Conceptualization: MI, EG, ANL

Funding acquisition: MI, EG, ANL

Data curation: MI

Methodology: MI, ANL

Visualization: MI, ANL

Investigation: MI, EG, ANL

Supervision: EG, ANL

Writing—original draft: MI

Writing—review & editing: MI, EG, ANL

## Data and materials availability

Code producing the data, analyses and figures we report on will be made available online upon publication. Individual language raw annotations and audio files can be retrieved from the corpora cited in Tables S1-S2.

## Supplementary Materials

**Table S1. The 44 studied languages (language corpora without IU annotations)**

Table S1 is available in an Open Science Foundation (OSF) repository.

**Table S2. The additional 4 languages studied for methods validation (language corpora with IU annotations)**

Table S2 is available in an <u>OSF repository</u>.

**Supplementary Text S1. Adding time-aligned annotations in English**

We relied on a corpus of spontaneous English speech, the Harvey Oral Narratives on Record (49). This corpus included only a segmentation into IUs but no time-stamped annotations at either word or IU level. To obtain the required timestamps we utilized and modified forced alignment tools put forth by Meta AI’s Massively Multilingual Speech project (79). Author MI manually segmented recordings to include the first ∼3:40 minutes, or continuing until the next speaker change. To create time-stamped annotations at the IU-levels for English, long audio files and text files containing a single IU per line were submitted to the forced alignment tool. The tool generates a *.json file consisting of the start and end times of each unit. In order to create time-stamped annotations at the word-level, we modified the alignment tool to create short audio files from each IU according to the identified start and end times, as well as a text file per IU containing a single orthographic word per line. Then, each of these shorter segments (audio and text file) were submitted to the forced alignment tool, generating a *.json file consisting of the start and end times of each word.

We processed the generated *.json files using custom-written scripts in R with the aid of the *rPraat* (80) and *dplyr* (73) packages. We created Praat (81) TextGrid files with annotations on both levels (IUs and words), and imported them into to ELAN (82). The time-stamped annotations were all manually checked and corrected by MI to exclude silent parts, breaths, and other noises that the forced alignment tool tended to include within annotations.

**Supplementary Text S2. Acoustic properties of automatically derived IUs reveal cross-linguistic audio-motor gestalts**

We segmented recordings of continuous, spontaneous speech in a wide variety of languages into IUs. We next turn to inspect, within each language, the resulting time- and scale-normalized envelope, f0 and harmonic ratio time courses of the automatically derived IUs. We assessed the statistical significance of the grand average time courses relative to 0 using a randomization procedure. Per language and per acoustic signal, we created 100 sets of surrogate average time courses. These surrogate time courses were calculated in a similar fashion (epochs were scaled and resampled, and then averaged per recording and across recordings per language), however on each iteration we circularly shifted each recording’s continuous acoustic signal (envelope, f0 and harmonic ratio) by a random number before epoching. Each iteration resulted in an average value per timepoint. However, we generated a single randomization distribution from this procedure by selecting, per iteration, the maximal absolute value across time. We then calculated for each timepoint whether the absolute observed value exceeds the 97.5% percentile of the randomization distribution of maximal statistics. This approach corrects for multiple comparisons over the different timepoints (76).

We find both similarities and differences in the acoustic realization of the detected IUs. Two characteristics appear across the board, in all languages: the speech envelope is consistently low at the very beginning of detected IUs (Fig. S1, left panel), and the harmonic ratio is consistently low at the very end of detected IUs (Fig. S1, right panel). Many languages also feature significantly low envelope values at the end of detected IUs and significantly low harmonic ratio values at the beginning. The harmonic ratio is low in silence by definition - given no speech there can be no voicing and therefore the calculated ratio between the periodic and aperiodic components of the signal is driven to 0. Therefore, the question arises of whether these deviations in envelope and harmonic ratio simply reflect silent pauses between IUs. However, the pattern of results suggests that the harmonic ratio is at least partly independent of the envelope and does not merely reflect silences: the envelope effect is consistent in each of the languages only at the beginning of detected IUs, whereas the harmonic ratio effect is consistent in each of the languages only at the end of detected IUs. The harmonic ratio might capture a different characteristic of IUs, the change in voice quality towards the ends of IUs that results in a low, raspy voice known as creaky voice (13). The vibration of the vocal folds tends to become irregular and particularly slow towards the end of IUs because of the gradual decrease in articulation pressure and the compression of the vocal folds.

In contrast to the trends in the envelope and harmonic ratio, deviations of f0 were less consistent in direction and modulation across the languages of our sample (Fig. S1, middle panel). The most stable pattern is a significant reduction in f0 towards the end of an IU, but this only occurs in 38 of the 48 languages (79%) and is distributed along the final third of the IU rather than being specific to the very end, as is the case for the harmonic ratio. In 3 languages, Cashinahua, Ruuli and Arapaho, the opposite pattern emerges (a significant increase in f0 at the end of the IU), and in 7 of the languages no significant pattern emerges towards the end of the IU.

**Fig. S1.**
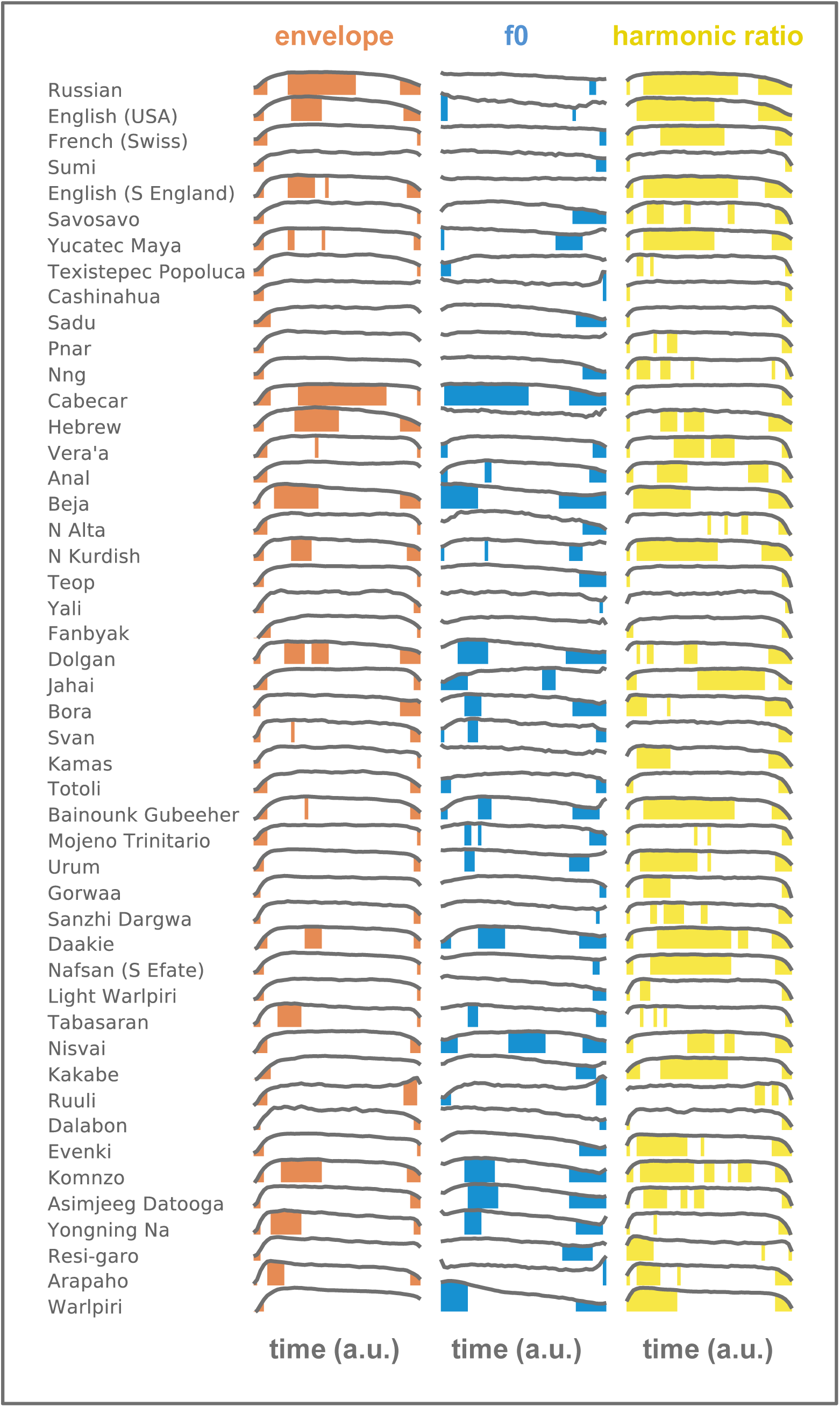
Envelope (left), f0 (middle) and harmonic ratio (right) grand average time courses of automatically derived IUs in 48 languages. Each gray line indicates a scale- and time-normalized grand average time course, computed by first averaging all normalized epochs within a recording, and then across the recordings in a given language. The y-axis range lies between [-1.5 1.5] z-scores and the x-axis range covers the entire epoch, from beginning to end. Colored regions under each curve correspond to portions of the time course that are significantly different (either greater or smaller) in comparison to surrogate time courses created in a randomization procedure, after correction for multiple comparisons across timepoints. The randomization procedure was designed to break the association between IU boundaries and a given acoustic signal by circularly shifting the acoustic signal.

